# Characterizing EEG Spectro-Temporal Variability Signatures in Alzheimer’s and Parkinson’s Disease

**DOI:** 10.64898/2026.03.07.710210

**Authors:** Yunier Prieur-Coloma, Pavel Prado, Wael El-Deredy, Alejandro Weinstein

**Affiliations:** Advanced Center for Electrical and Electronic Engineering, Universidad Técnica Federico Santa María, Valparaíso, Chile; Facultad de Ciencias de la Rehabilitación y Calidad de Vida, Universidad San Sebastián, Santiago de Chile, Chile; Escuela de Ingeniería Civil Biomédica, Universidad de Valparaíso, Valparaíso, Chile; Department of Electronic Engineering, Universidad Técnica Federico Santa María, Valparaíso, Chile

**Keywords:** electroencephalography, EEG spectral features, neurodegenerative diseases

## Abstract

We present an EEG-based approach to characterize disease-related spectro-temporal signatures in Alzheimer’s disease (AD) and Parkinson’s disease (PD). To this end, key spectral features were first identified using explainable machine learning, and their temporal dynamics were then examined to characterize variability patterns and statistical properties. EEG recordings were segmented into non-overlapping 4-s epochs, from which spectral features based on relative band power and spectral entropy were extracted. Random Forest classifiers were trained to discriminate individual subjects with AD and PD from healthy controls (HC) using a Leave-One-Subject-Out Cross-Validation (LOSOCV) strategy. The most discriminative spectral features and the directionality of their contributions were identified through a SHAP-based explainable analysis. Subsequently, the temporal dynamics of the key features were analyzed to characterize disease fingerprints in terms of variability at both inter-subject and intra-subject levels and their distributional profiles. Our results confirmed spectral slowing in both disorders and revealed disorder-specific differences in the dominant spectral markers: the theta/alpha ratio was the most influential feature for AD, whereas mean relative theta power was the primary feature for PD discrimination. We show that increased variability in key spectral features is a distinguishing signature of AD and PD, with disease groups exhibiting greater inter-subject heterogeneity and higher intra-subject temporal variability than HC. Moreover, the key features showed heavy-tailed behavior, for which a lognormal model provided a plausible fit across groups. We conclude that this EEG-based characterization provides a meaningful avenue for tracking deviations from healthy neural activity.

## I. Introduction

Neurodegenerative disorders represent a major global health problem due to their high prevalence and progressive nature [1]. Currently, more than 57 million people worldwide are affected by some form of neurodegenerative disease [2]. This number is expected to double every 20 years, mainly driven by population aging [3]. Among these conditions, Alzheimer’s disease (AD) and Parkinson’s disease (PD) are the most prevalent [4]. AD is the leading cause of dementia [5], while PD is one of the most common movement disorders [6]. Both diseases are characterized by progressive alterations of brain function that, at advanced stages, lead to cognitive, functional, and behavioral impairments. Therefore, non-invasive and low-cost tools are crucial for monitoring changes in neural activity toward disease conditions [7].

Electroencephalography (EEG) is a non-invasive technique that measures brain electrical activity with high temporal resolution [8]. It is sensitive to alterations in neural oscillatory dynamics and suitable for investigating neurodegenerative disorders [9], [10]. A growing body of literature has consistently reported disease-related alterations across canonical EEG frequency bands [11]–[14]. Recent works have combined EEG spectral features with machine learning techniques to discriminate between AD and PD [15]–[17]. Early work by Fiscon et al. analyzed AD and mild cognitive impairment (MCI) using frequency-domain features derived from fast Fourier transform and discrete wavelet transform coefficients across the canonical EEG frequency bands [18]. Using decision tree–based classifiers, the authors evaluated the discrimination of MCI and AD relative to healthy control (HC) subjects. The work proposed by Miltiadous et al. [19] used energy-based spectral features and statistical dispersion measures across the five canonical EEG bands to discriminate AD and frontotemporal dementia (FTD). Their study evaluated the performance of multiple classifiers, including decision trees, random forests, multilayer perceptrons, support vector machines (SVMs), Naïve Bayes, and k-nearest neighbors (k-NN). More recent studies have emphasized the combination of complementary EEG spectral features and inter-band power ratios to improve the AD classification performance [20]–[22]. In parallel, other studies have shown alterations in EEG spectral features associated with PD [23], [24]. Jaramillo-Jiménez et al. conducted a large multicenter study to characterize PD-related spectral changes [25]. Their results consistently showed increased theta power in patients with PD across four independent cohorts. In [26], band power, variance, and entropy-based spectral features were used to discriminate PD in on- and off-medication states from HC. Multiple classifiers were evaluated, including Random Forests, linear and quadratic discriminant analysis, SVMs, and k-NN. However, these studies were focused on optimizing classification performance, rather than uncovering disease-specific spectral signatures.

Other works have linked classifier decisions to physiologically meaningful EEG features, confirming well-established signatures of neurodegenerative disorders, including spectral slowing in AD and PD [27], [28]. Moreover, feature ranking strategies have highlighted discriminative spectral patterns with clear clinical relevance [29]. However, these approaches primarily rely on feature-importance representations, thereby overlooking the temporal organization and statistical structure of the discriminative features. As a result, potentially informative properties, such as the temporal dynamics of key features and their distributional profiles, remain uncharacterized. Addressing this gap may therefore provide additional clues about disease-related electrophysiological fingerprints. The hypothesis is that AD and PD are reflected not only in the magnitude of EEG spectral features but also in their temporal properties, such that the dynamics of these features encode disease fingerprints and may serve as markers for tracking shifts in brain activity toward disease conditions.

In this paper, an EEG-based approach is presented to characterize disease-related spectro-temporal signatures in AD and PD. Explainable machine learning is first used to identify the key spectral features for each condition, and their temporal dynamics are then characterized to reveal disease fingerprints encoded in feature variability. Finally, the statistical profiles of the identified features are modeled using lognormal distributions, providing an explicit distributional characterization. To the best of our knowledge, this is the first study to characterize the temporal dynamics and distributional profiles of key spectral features in AD and PD. Overall, the presented approach provides a comprehensive characterization of disease-related signatures in terms of relevance, variability, and distributional structure, opening new avenues for tracking deviations from healthy neural activity.

## II. Methods

In this work, an EEG-based approach is presented to characterize the temporal dynamics of key spectral features in AD and PD. As summarized in Fig. 1, the pipeline comprises five stages: (i) data selection from resting-state EEG recordings of AD, PD, and HC subjects; (ii) pre-processing and segmentation into non-overlapping epochs; (iii) extraction of spectral features per epoch; (iv) subject-level classification using a Random Forest model coupled with SHAP for interpretability; and (v) temporal and statistical characterization of identified key features to quantify variability and distributional profiles. The details of each stage are described in the following subsections.

**Fig. 1:**
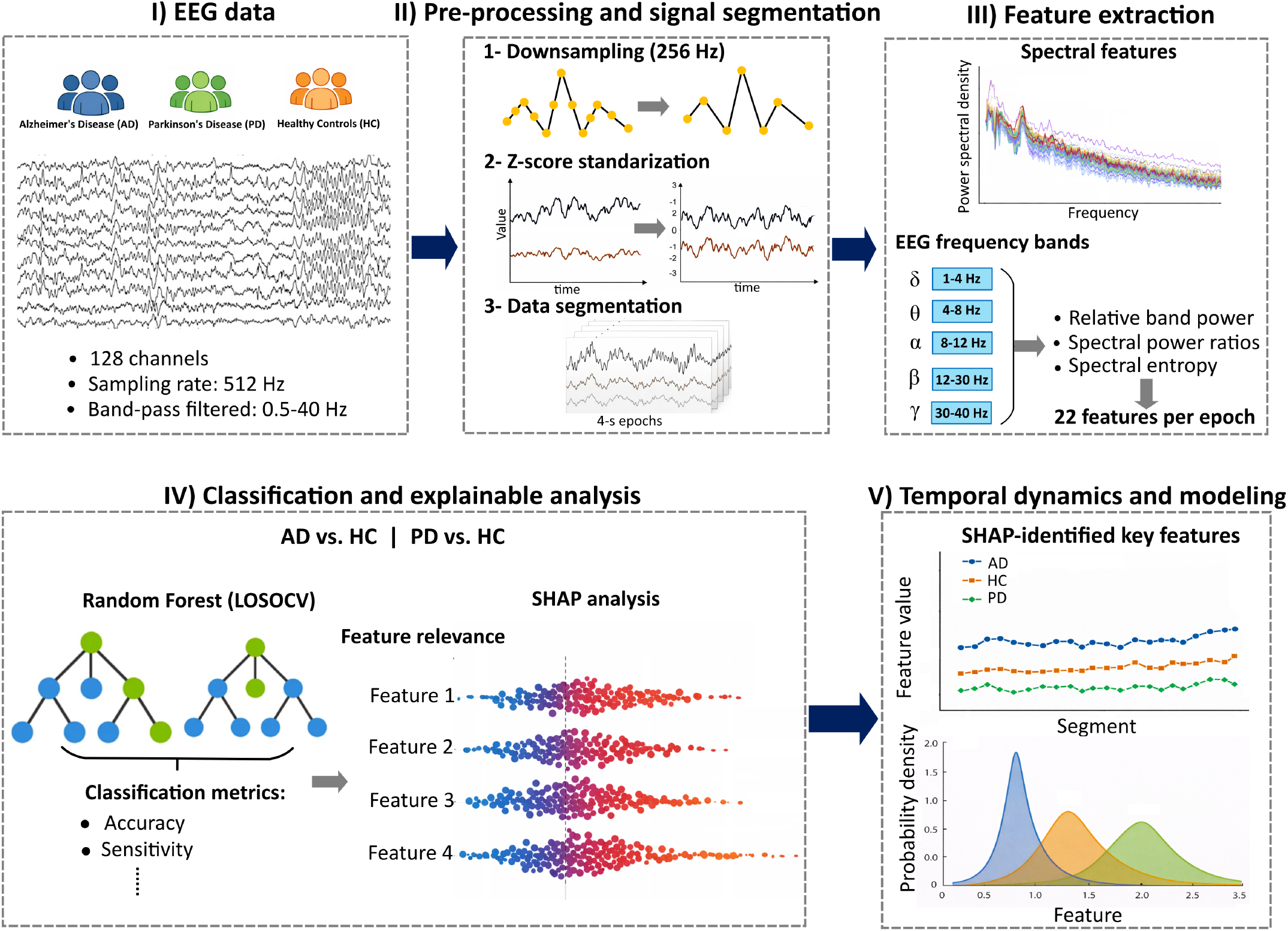
Overview of the proposed EEG-based approach. (I) Resting-state EEG recordings from Alzheimer’s disease (AD), Parkinson’s disease (PD), and healthy control (HC) subjects are used. (II) Pre-processing includes downsampling, z-score standardization, and segmentation into non-overlapping 4-s epochs. (III) Spectral features based on relative band power, spectral power ratios, and spectral entropy are extracted for each epoch. (IV) Random Forest classifiers are trained under a leave-one-subject-out cross-validation strategy, and model interpretability is achieved using SHAP analysis to identify the most discriminative spectral features. (V) Temporal dynamics and statistical modeling of key features are analyzed to characterize disease-related variability and deviations from healthy neural activity.

### A. Data Description

We used resting-state EEG recordings from a publicly available dataset [30]. The analyzed data comprise patients with AD and PD as well as HC subjects. Table I presents the clinical and demographic characteristics of each group.

**TABLE I:** Clinical-demographic characteristics of the study population.

| Characteristics | HC (n=12) | AD (n=19) | PD (n=22) |
| --- | --- | --- | --- |
| Age | 72.6 ± 4.2 | 77.9 ± 8.9 | 63.5 ± 7.9 |
| Females (%) | 50 | 57.8 | 54.5 |

The original dataset comprises resting-state EEG acquired using a 128-channel BioSemi ActiveTwo system equipped with pin-type active sintered Ag/AgCl electrodes, with participants resting in an eyes-closed condition. Each recording lasted approximately 10 minutes. According to the dataset description, the signals were re-referenced to the common average, band-pass filtered between 0.5 and 40 Hz, and resampled to 512 Hz. EEG artifacts associated with eye movements and blinking were corrected using Independent Component Analysis.

### B. Pre-processing and EEG Signals Segmentation

EEG recordings were downsampled from 512 Hz to 256 Hz to reduce computational complexity and storage requirements. All signals were then standardized using the z-score transformation (zero mean and unit variance) applied independently to each EEG channel within each subject. Machine learning–based classification systems typically require a sufficiently large number of samples to achieve robust generalization. In EEG-based disorder classification, a common strategy is to segment continuous recordings into short, informative epochs, which increases the effective sample size while preserving the intrinsic characteristics of the signal [31]–[33]. Following recent studies, EEG recordings from all conditions were segmented into non-overlapping 4-s epochs [34], [35]. This procedure resulted in 2413, 3565, and 1739 EEG segments for the AD, PD, and HC groups, respectively.

### C. Feature Extraction

Spectral features were extracted from each EEG segment based on statistical descriptors of relative band power and spectral entropy, which have been widely used in previous studies to discriminate AD and PD from HC [19], [26], [29], [29], [33].

Relative band power (RBP) quantifies the proportion of signal power contained within a specific frequency band relative to the total power across the analyzed frequency range. The power spectral density (PSD) was estimated with Welch’s method with a Hamming window [36], using 1-s segments with 50% overlap and a resulting frequency resolution of Δ*f* = 0.25 Hz. RBP was then computed for the five canonical EEG frequency bands: delta *δ* (1–4 Hz), theta *θ* (4–8 Hz), alpha *α* (8–12 Hz), beta *β* (12–30 Hz), and gamma *γ* (30–40 Hz) [37], as follows:

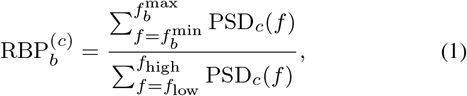

where PSD_*c*_(*f*) denotes the power spectral density at frequency *f* for channel *c*, 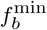 and 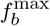 define the frequency limits of band *b*, and *f*_low_ = 1 Hz and *f*_high_ = 40 Hz correspond to the minimum and maximum frequencies considered in the analysis, respectively. To summarize the distribution of RBP across EEG channels, statistical descriptors were computed for each frequency band from the RBP values across channels. Specifically, the mean, median, and standard deviation of the RBP were calculated across channels, providing complementary information on the central tendency and spatial variability of oscillatory activity within each EEG frequency band.

Spectral power ratios (SPRs) were computed to characterize the relative balance between slow and fast oscillatory components. SPRs were derived from the mean RBP values across EEG channels. For each EEG segment, a spectral power ratio was defined as follows:

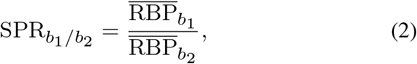

where *b*_1_ and *b*_2_ denote two distinct frequency bands and 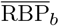 represents the mean relative band power across EEG channels for band *b*. The following spectral ratios were considered: *δ/θ, δ/α, δ/β, θ/α, θ/β*, and *α/β*.

In addition to band-specific features, wide-band spectral entropy (SE) was computed across the full frequency range (1–40 Hz) to provide a global measure of spectral complexity. Spectral entropy quantifies the degree of irregularity and dispersion of spectral power across frequencies. SE was computed from the PSD averaged across EEG channels as follows:

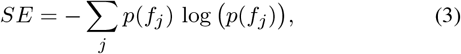

where *p*(*f*_*j*_) denotes the normalized channel-averaged PSD at the *j*-th frequency bin, defined as:

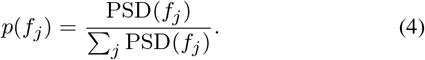

In total, 22 spectral features were extracted from each EEG segment. These comprised: (i) statistical descriptors (mean, median, and standard deviation) of RBP computed for each canonical frequency band; (ii) inter-band spectral power ratios computed from the mean relative band power; and (iii) wide-band spectral entropy.

### D. Classification Model and Explainable Analysis

Two classification problems were addressed: AD vs. HC and PD vs. HC. For both cases, a Random Forest (RF) classifier was employed, given its robust performance in EEG-based classification studies [19], [33]. RF is an ensemble learning method that constructs multiple decision trees from bootstrap samples of the training data and combines their outputs through majority voting, thereby reducing variance and improving generalization.

The classifier was trained and validated using spectral features extracted from multiple EEG segments. Each segment from a given subject represents one sample for the model. However, these samples are not independent (several segments are from the same subject). If segments from the same subject are simultaneously included in both the training and testing sets (data leakage), the model’s performance will be artificially inflated. To avoid this issue and ensure strict subject-level separation, a Leave-One-Subject-Out Cross-Validation (LOSOCV) strategy was adopted [38]. In LOSOCV, all segments from a single subject are held out at each iteration, while the classifier is trained on the remaining subjects’ data. The trained model is then evaluated on the left-out subject’s segments. This procedure is repeated for each subject, yielding an unbiased estimation of generalization performance. Note that model predictions are generated at the segment level, whereas the final decision must be made at the subject level. Following the approach proposed by Sibilano et al. [32], subject-level predictions were obtained by averaging the predicted class probabilities across all segments belonging to the same subject. The final class label was assigned according to the class with the highest mean probability. Model performance was evaluated at the subject level using Accuracy, Sensitivity, Specificity, Precision, and F1-score (see Supplementary Material A).

RF classifiers provide out-of-bag (OOB) feature importance estimates derived from prediction errors on samples not used during tree training. These importance scores have been used to identify relevant physiological markers [39]. However, OOB feature importance captures only global feature relevance. It does not explain how features contribute to individual predictions, nor does it indicate the directionality of their effects. To overcome these limitations and improve interpretability, RF classification was complemented with a SHAP-based explainable analysis.

We used TreeSHAP, an efficient SHAP implementation specifically designed for tree-based models [40]. SHAP provides a unified game-theoretic framework for quantifying each feature’s contribution to an individual prediction. TreeSHAP computes Shapley values for ensemble tree models by leveraging the internal structure of the ensemble. A Shapley value represents the marginal contribution of a feature to a given prediction, considering all possible coalitions of features. Positive SHAP values indicate that a feature pushes the model output toward the positive class (AD or PD), whereas negative values push the prediction toward the control class (HC). This enables not only the ranking of the most influential features but also the explicit characterization of the direction and magnitude of their effects at the group level. Using TreeSHAP thus enables a fine-grained, disorder-specific interpretability analysis that complements global model performance metrics and offers neurophysiologically meaningful insights into the spectral features associated with AD and PD [28].

### E. Temporal Dynamics and Modeling of Discriminative Spectral Features

Temporal dynamics of the most discriminative spectral features identified by the SHAP analysis were obtained by concatenating segment values across time. The group-level temporal profiles were summarized using the mean and standard deviation across subjects at each segment. Statistical analyses were conducted to assess group-level differences in temporal variability. Intra-subject variability was used as a measure of signal stability and was quantified for each subject by computing the interquartile range (IQR) of feature values across EEG segments. Group differences in intra-subject variability between disease conditions (AD and PD) and HC were first evaluated using the Mann-Whitney U test.

To test for potential age confounding, group differences in variability were further assessed using analyses of covariance (ANCOVA) [41]. Group was defined as the independent variable, Age was included as a covariate, and the IQR of each key feature computed across EEG segments was used as the dependent variable. The slope homogeneity was assessed by including a Group×Age interaction term. For each term (Group, Age, and Group×Age), the *F* statistic was computed, and statistical significance was assessed using the associated *p*-value. Partial eta squared 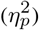 was calculated as a measure of effect size. For these analyses, IQR values were log-transformed to reduce distributional asymmetry.

To characterize the statistical properties of key spectral features across disease and healthy conditions, a distributional analysis was conducted. The empirical distributions were examined for AD, PD, and HC groups. Following the methodology proposed by Clauset et al. [42], the data were fitted with power-law, lognormal, and exponential models. Model fitting was performed using maximum likelihood estimation and restricted to values *x* ≥ *x*_min_, ensuring that all candidate distributions were compared over the same support (see Supplementary Material C). The lower bound *x*_min_ was estimated following the approach described in [43]. The optimal value 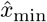 was selected as the one that minimizes the distances between the empirical distribution and the fitted model for *x* ≥ *x*_min_.

The Kolmogorov–Smirnov distance was used as a nonparametric measure, quantifying the maximum distance between the cumulative distribution functions (CDFs) of the empirical data and the fitted model, as follows:

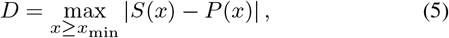

where *S*(*x*) denotes the empirical CDF of the data restricted to values *x* ≥ *x*_min_, and *P* (*x*) denotes the CDF of the fitted model over the same range. The estimate 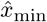 corresponds to the value of *x*_min_ that minimizes *D*.

Model selection was conducted using log-likelihood ratio (*R*) tests. Given two candidate models ℳ_1_ and ℳ_2_, the log-likelihood ratio was defined as follows:

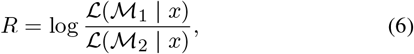

where ℒ (ℳ_*i*_ | *x*) denotes the likelihood of model ℳ_*i*_ given the data. Positive values of *R* indicate a preference for model ℳ_1_ over ℳ_2_, while negative values indicate the opposite. Statistical significance was assessed using the associated *p*-value, as described in [42].

All code required to reproduce the analyses in this study is publicly available at Zenodo [44].

## III. Results

In this section, we report the classification performance for AD and PD discrimination using the extracted spectral features and the RF classifier evaluated under the LOSOCV strategy. Next, we summarize the SHAP-based explainable analysis, highlighting the most influential features and the directionality of their contributions. Finally, we report the results of the temporal dynamics analysis and distributional modeling of key features across all conditions.

### A. Classification and Explainable Analysis

Table II presents the classification performance for both AD vs. HC and PD vs. HC. In the AD vs. HC scenario, the classifier achieved an accuracy of 67.7%, indicating a moderate discrimination capability. Sensitivity was 73.6%, suggesting that most AD subjects were correctly identified; however, the lower specificity (58.3%) suggests a higher rate of false positives among HC subjects. In the PD vs. HC case, performance was stronger, with an accuracy of 82.2% and a sensitivity of 95.5%, highlighting the model’s high ability to detect PD, while specificity remained at 58.3%. The precision obtained for AD and PD was 73.6% and 80.3%, respectively, with corresponding F1-scores of 73.6% and 87.5%, indicating that positive predictions were robust, particularly for PD.

**TABLE II:** Classification performance for AD vs. HC and PD vs. HC using a Random Forest classifier under the LOSOCV strategy. Accuracy (Acc), sensitivity (Sen), specificity (Spe), precision (Prec), and F1-score (F1).

| Case | Acc<br>(%) | Sen<br>(%) | Spe<br>(%) | Prec<br>(%) | F1<br>(%) |
| --- | --- | --- | --- | --- | --- |
| AD vs. HC | 67.7 | 73.6 | 58.3 | 73.6 | 73.6 |
| PD vs. HC | 82.5 | 95.4 | 58.3 | 80.7 | 87.5 |

Fig. 2 presents the SHAP-based feature importance for the positive classes in both classification tasks (AD and PD). Both plots jointly show the relative contribution of the 22 extracted spectral features, ranked from highest to lowest relevance, and their directional influence on the model’s predictions. As shown in Fig. 2(a), for the AD prediction task, the most influential feature was the *θ/α* ratio. Fig. 2(b) shows that, for PD detection, the mean relative *θ* power emerged as the most relevant feature. As in the AD case, the *θ/α* ratio ranked among the top contributing features, suggesting that this SPR plays a key discriminative role across conditions.

**Fig. 2:**
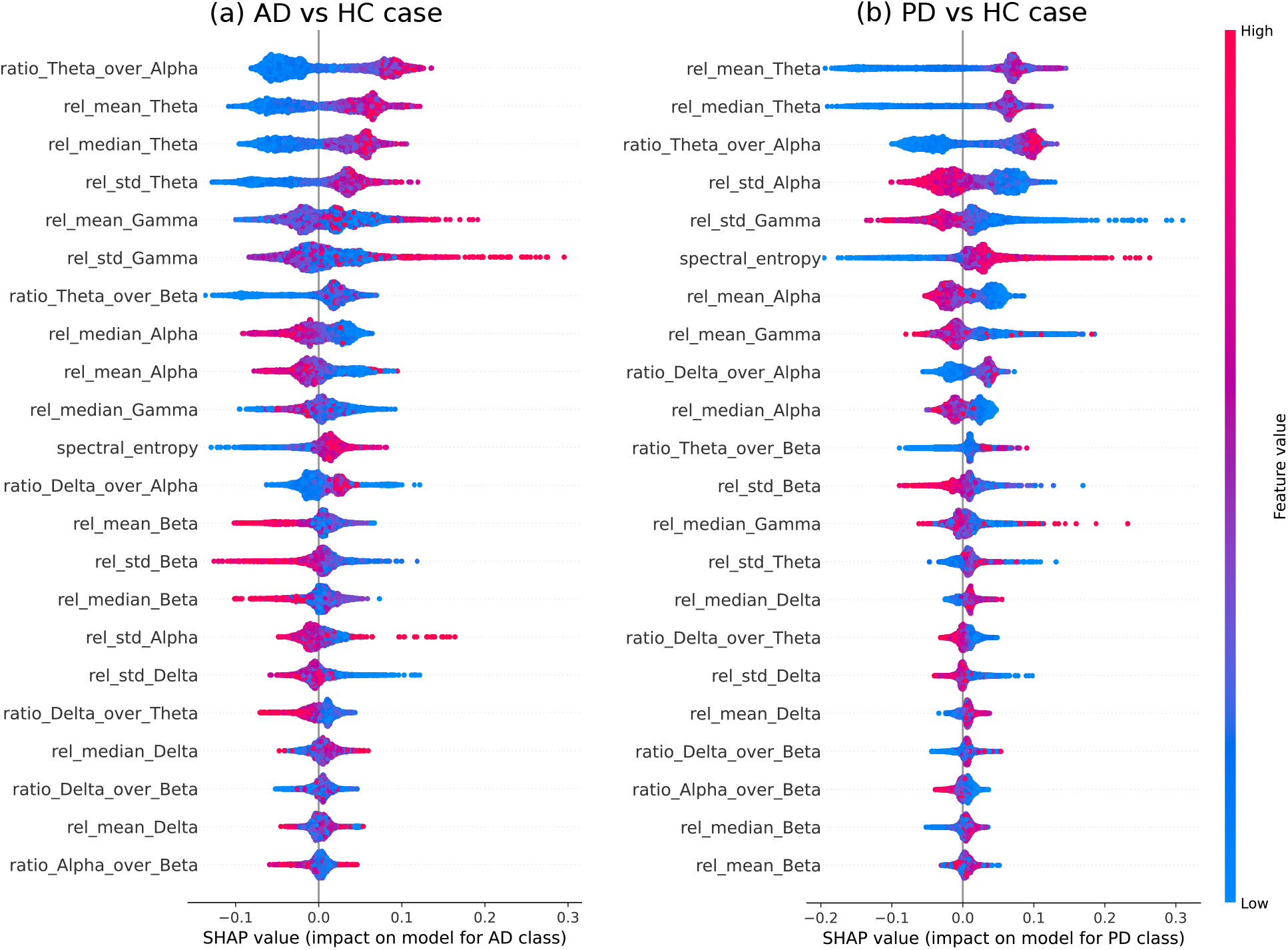
SHAP-based feature importance for the two classification tasks, (a) corresponds to the AD vs. HC classification and panel (b) to the PD vs. HC classification. Each plot displays the contribution and direction of influence of the 22 spectral features, ordered from highest to lowest relevance for the respective positive class.

The mean relative *θ* power and the *θ/α* ratio consistently rank among the top three most influential features for AD and PD classification (see Fig. 2).

### B. Temporal Dynamics and Modeling

As shown in Section III-A, the *θ/α* ratio and the mean relative *θ* power emerged as the most relevant spectral features for discriminating AD and PD from HC (see Fig. 2). Fig. 3 presents the temporal dynamics of these features for the AD vs. HC and PD vs. HC cases, where solid lines and shaded areas denote the mean and ±1 standard deviation across subjects for each segment, respectively. Specifically, Figs. 3(a) and 3(b) show the *θ/α* ratio and mean relative *θ* power for AD vs. HC, respectively, whereas Figs. 3(c) and 3(d) show the corresponding results for PD vs. HC. Across both comparisons, disease groups exhibited high inter-subject variability, reflected by wide ±1 standard deviation bands across all EEG segments (Figs. 3(a)–(d)). This pattern was more pronounced for the *θ/α* ratio than for mean relative *θ* power. In contrast, such variability was not observed for SHAP low-relevance features (see Supplementary Material B). In addition, as shown in Fig. 4, temporal variability differed significantly between disease conditions and healthy controls for both features. For the *θ/α* ratio, differences were significant (AD vs. HC: *p <* 0.01; PD vs. HC: *p <* 0.001), for the mean relative *θ* power, significant differences were also found (AD vs. HC: *p <* 0.05; PD vs. HC: *p <* 0.001; Mann-Whitney U test). As reported in Table III, the main effect of Group remained significant after age adjustment for both the *θ/α* ratio (*p* = 3.70 × 10^*−*4^) and mean relative *θ* power (*p* = 1.07 × 10^*−*3^). For the *θ/α* ratio, the effect of Age was not significant (*p* = 0.193), whereas for mean relative *θ* power a significant age effect was observed (*p* = 0.023). The Group×Age interaction was not significant for either feature (both *p* ≥ 0.29), indicating comparable age-variability slopes across groups.

**Fig. 3:**
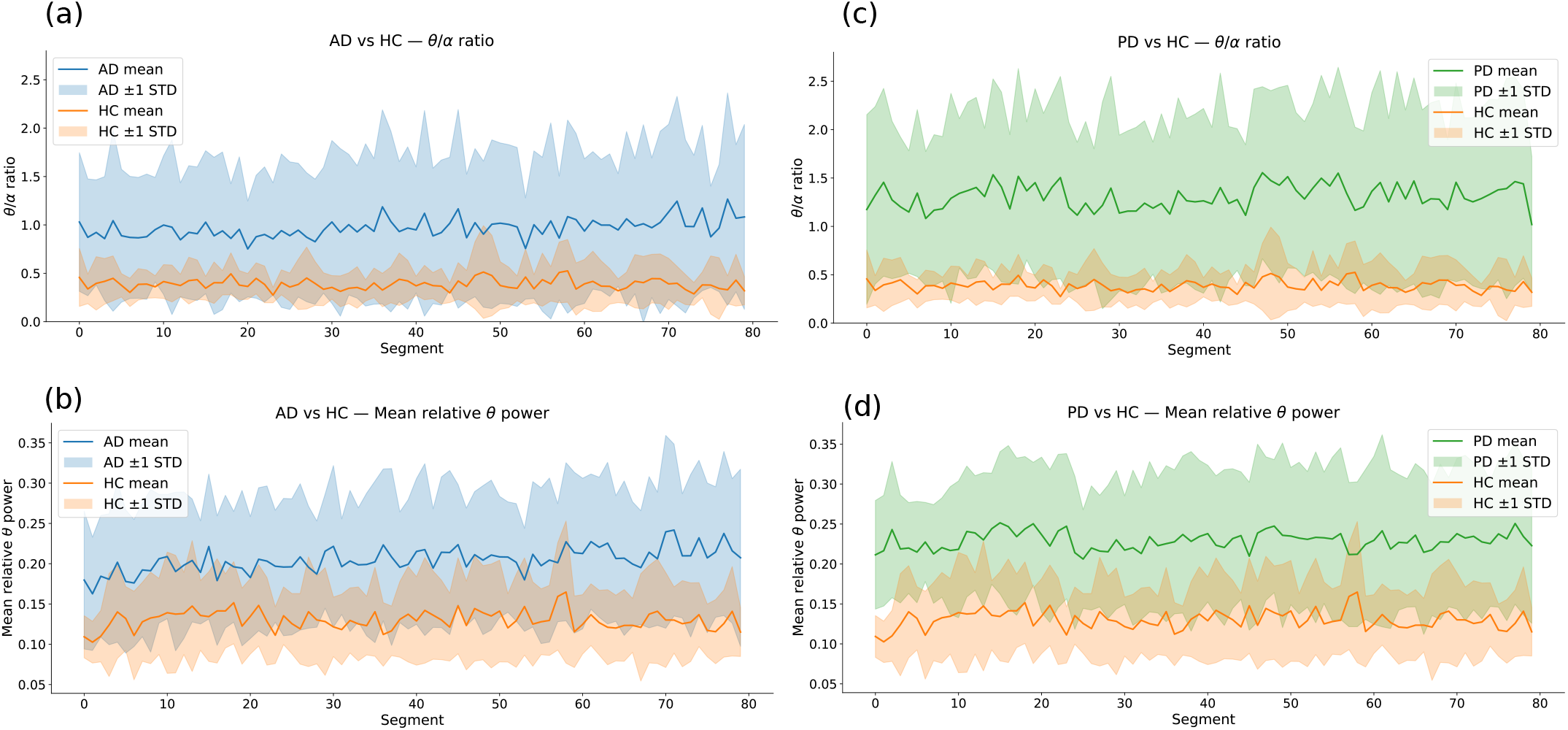
Temporal dynamics of key spectral features for AD vs. HC and PD vs. HC. The *θ/α* ratio and mean relative *θ* power are shown across EEG segments. Panels (a)–(b) correspond to AD vs. HC comparisons, whereas panels (c)–(d) depict PD vs. HC comparisons. Solid lines denote the mean temporal dynamics for each condition, and shaded regions represent ±1 standard deviation across subjects for each segment.

**Fig. 4:**
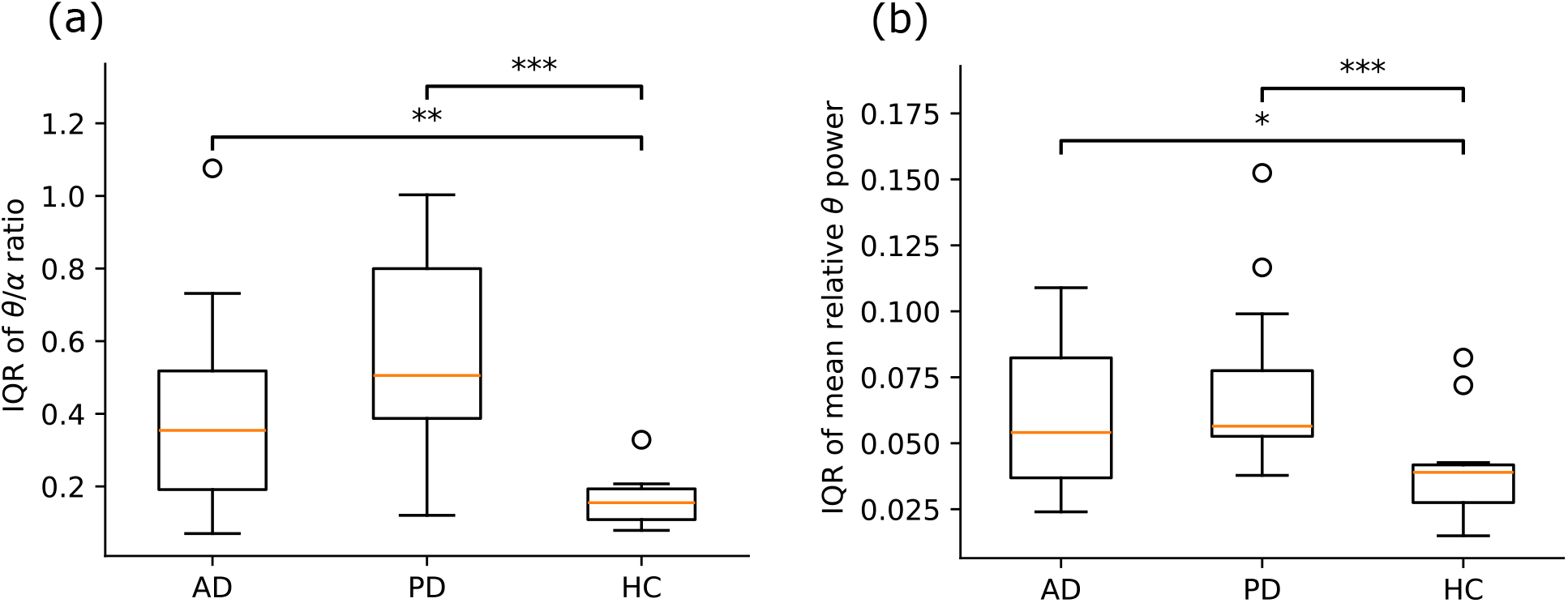
Intra-subject temporal variability of key spectral features across groups. (a) Variability of the *θ/α* ratio and (b) variability of mean relative *θ* power. For each subject, variability was quantified using the within-subject interquartile range (IQR) computed across EEG segments. Group differences were assessed using the Mann–Whitney U test (\**p <* 0.05, \*\**p <* 0.01, \*\*\**p <* 0.001).

**TABLE III:** ANCOVA results testing the main effects of Group and Age, and their interaction, on intra-subject variability.

| | $\theta/\alpha$ ratio (IQR) | | | |
| --- | --- | --- | --- | --- |
| | $df$ | $F$ | $p$ | $\eta_p^2$ |
| Group | **2, 47** | **9.39** | **< 0.001** | **0.28** |
| Age | 1, 47 | 1.74 | 0.192 | 0.03 |
| Group×Age | 2, 47 | 0.85 | 0.43 | 0.03 |
| | Mean relative $\theta$ power (IQR) | | | |
| | $df$ | $F$ | $p$ | $\eta_p^2$ |
| Group | **2, 47** | **7.94** | **< 0.01** | **0.25** |
| Age | **1, 47** | **5.48** | **< 0.05** | **0.10** |
| Group×Age | 2, 47 | 1.26 | 0.29 | 0.05 |

Table IV presents the model-comparison results for the distributional analysis described in Section II-E. Although no single distribution consistently outperformed all alternatives across features and conditions, the lognormal model provided the most consistent fit, generally outperforming the power-law model and yielding comparable performance to the exponential model across both disease and healthy groups. Accordingly, the lognormal distribution was selected as a common reference model for subsequent between-group comparisons.

**TABLE IV:**
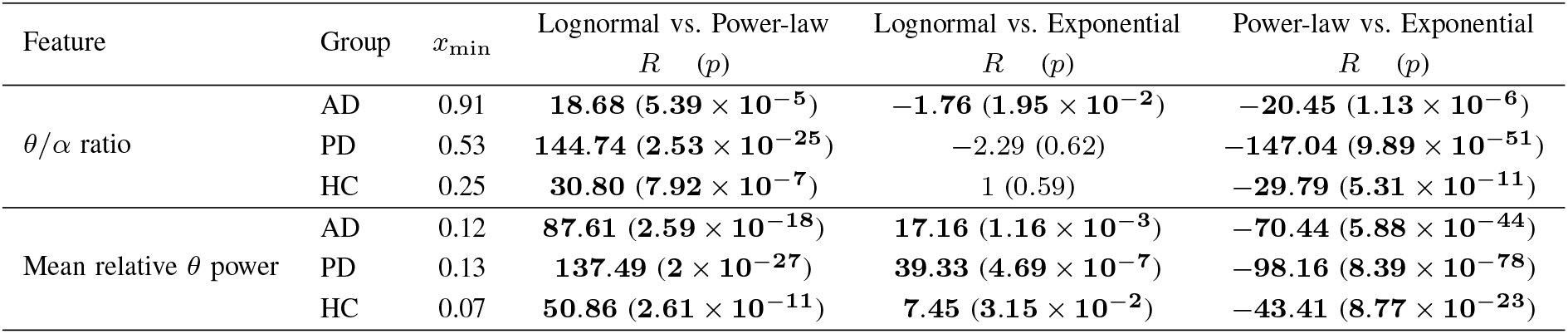
Log-likelihood ratio (*R*) and associated statistical significance (*p*) for model comparisons of the *θ/α* ratio and mean relative *θ* power.

| Feature | Group | $x_{\min}$ | Lognormal vs. Power-law | | Lognormal vs. Exponential | | Power-law vs. Exponential | |
| --- | --- | --- | --- | --- | --- | --- | --- | --- |
| | | | $R$ | ($p$) | $R$ | ($p$) | $R$ | ($p$) |
| $\theta/\alpha$ ratio | AD | 0.91 | **18.68** | ($5.39 \times 10^{-5}$) | **-1.76** | ($1.95 \times 10^{-2}$) | **-20.45** | ($1.13 \times 10^{-6}$) |
| | PD | 0.53 | **144.74** | ($2.53 \times 10^{-25}$) | -2.29 | (0.62) | **-147.04** | ($9.89 \times 10^{-51}$) |
| | HC | 0.25 | **30.80** | ($7.92 \times 10^{-7}$) | 1 | (0.59) | **-29.79** | ($5.31 \times 10^{-11}$) |
| Mean relative $\theta$ power | AD | 0.12 | **87.61** | ($2.59 \times 10^{-18}$) | **17.16** | ($1.16 \times 10^{-3}$) | **-70.44** | ($5.88 \times 10^{-44}$) |
| | PD | 0.13 | **137.49** | ($2 \times 10^{-27}$) | **39.33** | ($4.69 \times 10^{-7}$) | **-98.16** | ($8.39 \times 10^{-78}$) |
| | HC | 0.07 | **50.86** | ($2.61 \times 10^{-11}$) | **7.45** | ($3.15 \times 10^{-2}$) | **-43.41** | ($8.77 \times 10^{-23}$) |

Fig. 5 shows the lognormal probability density functions estimated for the *θ/α* ratio and the mean relative *θ* power in AD, PD, and HC. For both features, AD and PD exhibit broader distributions with heavier right tails than HC. For the *θ/α* ratio, this effect is more pronounced in the disease groups, with greater variability in AD (*σ* = 0.82) and PD (*σ* = 0.76) than in HC (*σ* = 0.51). For the mean relative *θ* power, the differences in variability are more subtle, with AD (*σ* = 0.44) and PD (*σ* = 0.37) relative to HC (*σ* = 0.37). The fitted lognormal parameters for each condition are reported in Supplementary Material C.

**Fig. 5:**
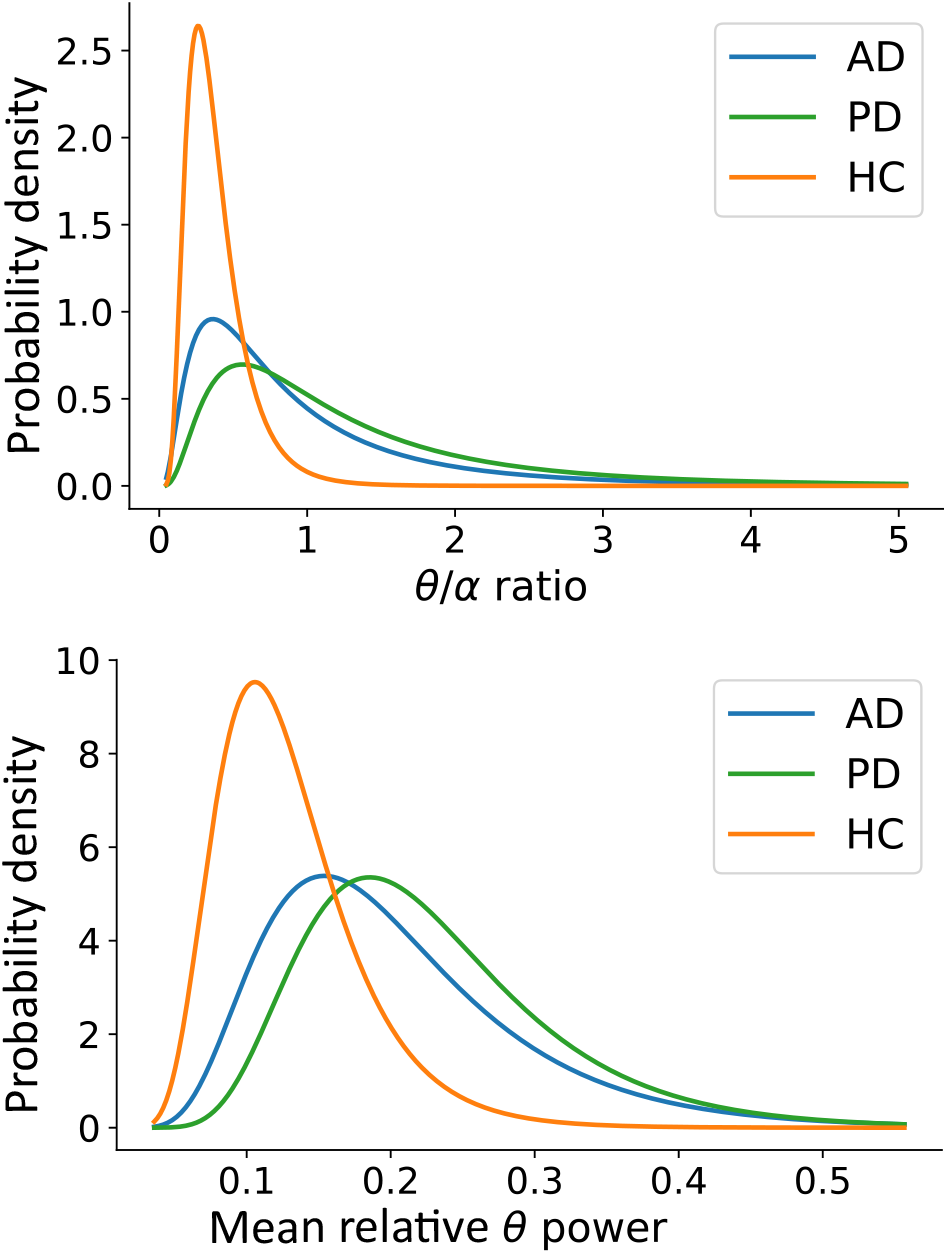
Estimated probability density functions of the lognormal model fitted by maximum likelihood for the *θ/α* ratio (top) and the mean relative *θ* power (bottom) in Alzheimer’s disease (AD), Parkinson’s disease (PD), and healthy control (HC) subjects. AD and PD subjects exhibit broader distributions and heavier right tails than healthy controls, indicating greater dispersion and a higher probability of extreme values.

## IV. Discussion

In this study, we present an EEG-based approach to characterize how the temporal dynamics of spectral features differ across AD, PD, and HC. Continuous resting-state EEG recordings were segmented into 4-s epochs, from which 22 spectral features were extracted across five EEG frequency bands. These features were used to address two classification problems (AD vs. HC and PD vs. HC), using an RF model evaluated with a LOSOCV strategy. The RF classifier was coupled with SHAP to identify the most relevant spectral features contributing to disease discrimination and to explicitly characterize their contribution. Finally, the presented approach extends beyond feature importance by examining the temporal dynamics and statistical properties of key features, thereby enabling their characterization in terms of variability and distributional profiles.

We achieved moderate accuracy for both AD and PD discrimination tasks (see Table II). The model exhibited a consistent performance pattern, with higher sensitivity than specificity. The achieved accuracies are comparable to those reported in prior EEG-based studies on AD and PD discrimination [45], [46]. It is worth noting that this work focuses on achieving a balanced trade-off between classification performance and model interpretability, with the aim of characterizing meaningful spectral signatures associated with each disorder.

Previous studies have consistently identified spectral slowing as a hallmark electrophysiological feature of AD and PD, characterized by increased power in low-frequency EEG bands (e.g., theta) and reduced power in higher-frequency oscillations (e.g., alpha and beta) [27], [47]. Our results provide a more fine-grained, disorder-specific characterization of spectral slowing. As shown in Fig. 2, the *θ/α* ratio was the most influential feature driving AD discrimination, whereas the mean relative *θ* power emerged as a dominant feature in the PD classification task. The SHAP-based explainable analysis revealed the directionality in these effects: higher *θ/α* values pushed the model toward the AD class, while increased mean relative *θ* power favored predictions toward PD. In contrast, higher relative power in faster frequency bands (alpha and beta) shifted predictions toward HC across both classification tasks. These results align with studies indicating that preserved fast-frequency activity is inversely related to neurodegenerative disease [48], [49].

It is widely accepted that EEG spectral signatures of AD and PD are primarily driven by changes in oscillatory activity [50]. By accounting for previously overlooked temporal dynamics of key EEG spectral features, variability patterns were examined at both inter- and intra-subject levels. Our results highlight increased variability at both levels. At the inter-subject level (Fig. 3), disease groups exhibited greater heterogeneity, with wider dispersion of feature values across individuals. This finding is consistent with recent evidence indicating that neurodegenerative diseases, including AD and PD, exhibit considerable spectral deviations within patients [51]. At the intra-subject level, temporal variability was significantly higher in AD and PD than in HC for both features (see Fig. 4), which is in line with previous evidence of aberrant neural synchrony in neurodegenerative disorders [52]. These differences were supported by unadjusted comparisons and were subsequently corroborated after age adjustment (see Table III), in which a significant main effect of Group was retained. These results indicate that AD and PD are associated with increased temporal variability beyond shifts in average spectral power. Our results suggest that increased temporal variability of the *θ/α* ratio reflects a dynamic biomarker sensitive to short-timescale fluctuations and may be suitable for monitoring subtle changes in brain function during early disease stages.

Finally, we provided an explicit distributional characterization of key spectral features across AD, PD, and HC subjects. In this context, lognormal distributions estimated from HC subjects may serve as a normative reference that reflects intrinsic physiological variability while preserving the characteristic spectral profile of a healthy brain. This modeling approach aligns with evidence that neural and physiological brain variables are inherently non-Gaussian, exhibiting skewed distributions with heavy tails and asymmetric fluctuations [53]. Within this framework, deviations from the control distributions can be used to identify subjects whose spectral profiles progressively diverge from normal patterns, potentially enabling the detection of disease-related alterations at early stages.

A limitation of this work is the relatively small sample size. Future work will focus on extending the proposed analysis to larger, multicenter datasets, incorporating recordings acquired across independent sites and protocols. This will enable a more reliable estimation of feature distributions and a more robust characterization of the statistical properties of EEG activity in healthy and disease-related conditions.

## V. Conclusion

In this work, an EEG-based approach for characterizing disease-related spectro-temporal signatures in AD and PD is presented. By integrating spectral feature extraction with explainable machine learning, the proposed approach enables the characterization of EEG alterations in terms of feature relevance, directionality, temporal dynamics, and variability. Our results indicate that AD and PD are associated with spectral slowing and increased variability in key features, characterized by larger inter-subject heterogeneity and intra-subject temporal variability. Finally, this study provides meaningful EEG markers and represents a promising tool for monitoring deviations from healthy neural activity in neurodegenerative disorders.

