## Supplementary Material for "Characterizing EEG Spectro-Temporal Variability Signatures in Alzheimer’s and Parkinson’s Disease"

Yunier Prieur-Coloma[1], Pavel Prado[2], Wael El-Deredy[3] and Alejandro Weinstein[4]

*[1]Advanced Center for Electrical and Electronic Engineering, Universidad Técnica Federico Santa María, Valparaíso, Chile.*

*[2]Facultad de Ciencias de la Rehabilitación y Calidad de Vida, Universidad San Sebastián, Santiago de Chile, Chile.*

*[3]Escuela de Ingeniería Civil Biomédica, Universidad de Valparaíso, Valparaíso, Chile.*

*[4]Department of Electronic Engineering, Universidad Técnica Federico Santa Maria, Valparaíso, Chile.*

#### A. Classification metrics

Model performance at the subject level was evaluated using Accuracy (Acc), Sensitivity (Sen), Specificity (Spe), Precision (Prec), and F1-score, all computed from the confusion matrix. For each binary classification scenario (e.g., AD vs. HC or PD vs. HC), the positive class corresponds to the disease group (AD or PD, respectively), and the negative class corresponds to HC. Let TP denote true positives, TN true negatives, FP false positives, and FN false negatives. The classification metrics are defined as:

$$Acc = \frac{TP + TN}{TP + TN + FP + FN}$$

$$Sen = \frac{TP}{TP + FN}$$

$$Spe = \frac{TN}{TN + FP}$$

$$Prec = \frac{TP}{TP + FP}$$

$$F1 = 2\frac{Prec \cdot Sen}{Prec + Sen}$$

### B. Temporal dynamics of low-relevance features identified by SHAP

Key features identified through the SHAP analysis exhibited pronounced variability at the inter-subject level within the disease groups. In contrast, this variability pattern was not observed for less relevant features. Supplementary Figures S1 and S2 illustrate the temporal dynamics of mean relative δ power, which emerged as one of the least relevant characteristics for discriminating AD and PD from healthy controls in the corresponding classification tasks.

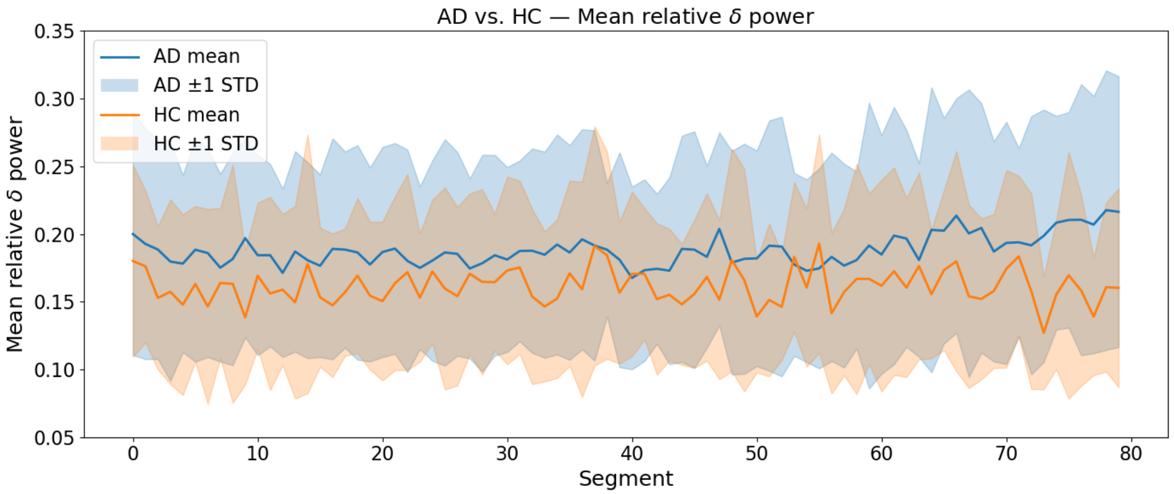


Supplementary Figure S1. Temporal dynamics of mean relative δ power for the AD vs. HC comparison. Solid lines show the group mean across segments, and shaded regions indicate ±1 standard deviation across subjects (inter-subject variability) at each segment. Mean relative δ power was ranked as a low-relevance feature in the SHAP-based analysis for AD discrimination.

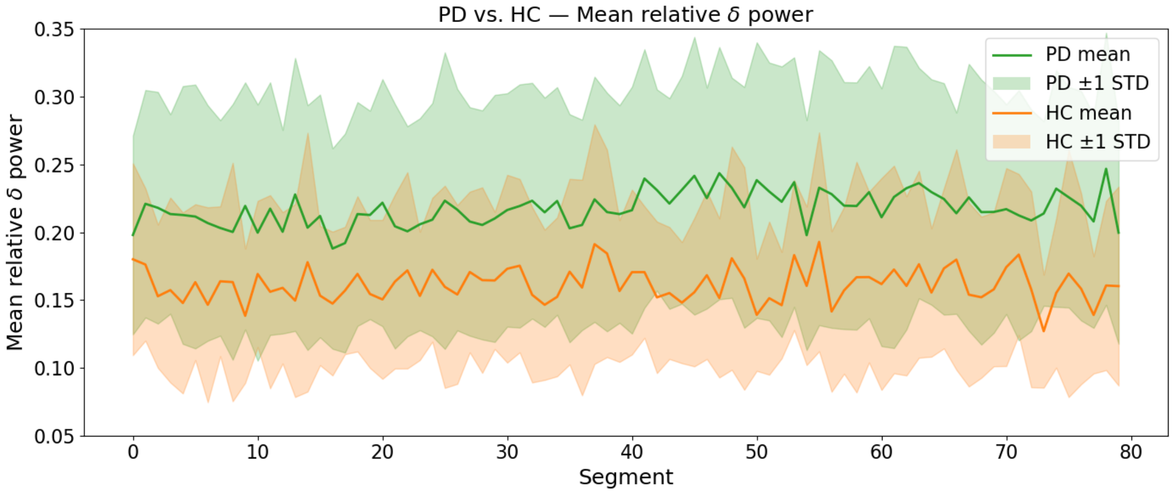


Supplementary Figure S2. Temporal dynamics of mean relative δ power for the AD vs. HC comparison. Solid lines show the group mean across segments, and shaded regions indicate ±1 standard deviation across subjects (inter-subject variability) at each segment. Mean relative δ power was ranked as a low-relevance feature in the SHAP-based analysis for AD discrimination.

#### C. Modeling of key features

Supplementary Figures S3 and S4 show the histograms of the two key features identified through SHAP analysis. A right-skewed pattern is observed across all groups (AD, PD, and HC).

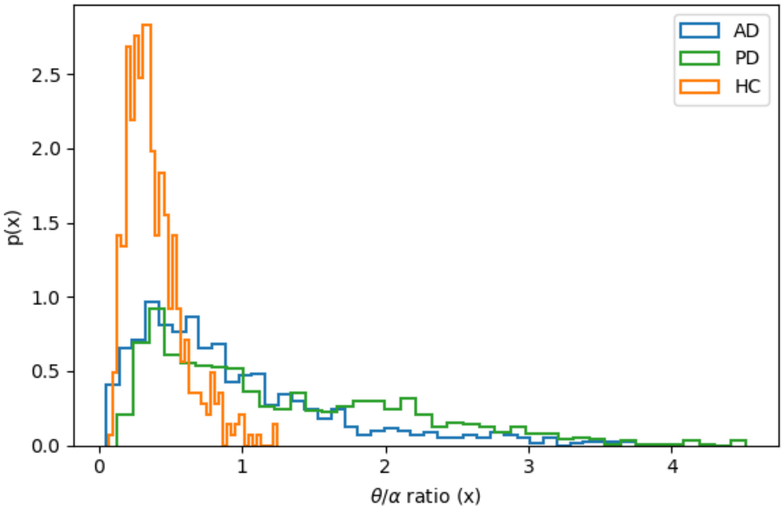


Supplementary Figure S3. Histogram of the $\theta/\alpha$ ratio feature for AD, PD and HC conditions.

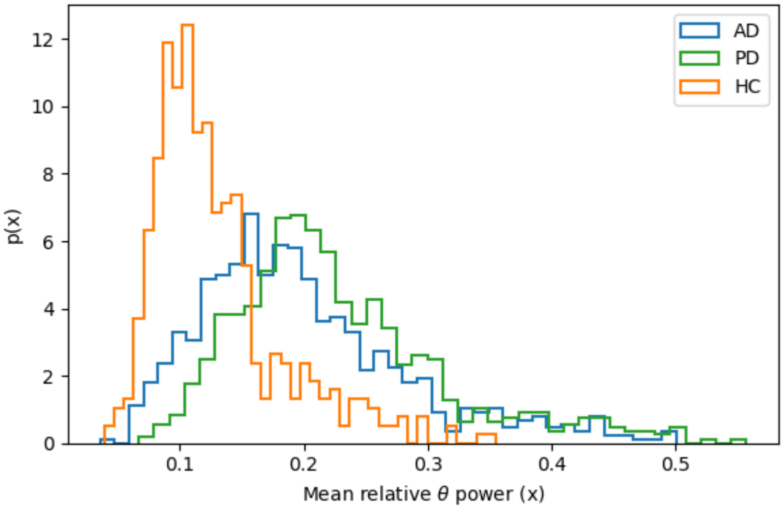


Supplementary Figure S4. Histogram of the mean relative $\theta$ power feature for AD, PD, and HC conditions.

However, histogram-based visualization is sensitive to the choice of the number of bins. To address this limitation, we further characterized the empirical distributions by fitting three candidate models: power-law, lognormal, and exponential.

As an illustrative example, consider testing whether a power-law is a plausible hypothesis for the data. A random variable $x$ is said to follow a power-law if it is drawn form a probability distribution:

$$p(x) \propto x^{-\alpha}$$

where $\alpha$ is the scaling parameter. In practice, power-law behavior typically holds only above some minimum value. For example, a continuous power-law distribution can be written as:

$$p(x) = Cx^{-\alpha}$$

where $C$ is a normalization constant. Assuming $\alpha > 1$, the density diverges as $x \to 0$, so this expression cannot hold for all $x \geq 0$. Therefore, a lower bound ($x_{min}$ is introduced, such that the power-law behavior applies for $x \geq x_{min}$.

Before estimating the scaling parameter $\alpha$, it is necessary to estimate $x_{min}$ in order to discard all samples below this lower bound, leaving only those values for which the power-law model provides a valid approximation. Here, we estimated $x_{min}$ by selecting the value that makes the empirical distribution of the data and the best-fitting power-law model as similar as possible for $x \geq x_{min}$.

To ensure a consistent basis for model comparison, the three candidate models (power-law, lognormal, and exponential) were fitted using the same estimated lower bound $x_{min}$ (i.e., all fits were performed on data with $x \geq x_{min}$).

Formally, the tail probability density functions of the three candidate models are:

Power-law:

$$p_{pow}(x) = \frac{\alpha - 1}{x_{min}} \left(\frac{x}{x_{min}}\right)^{-\alpha}, \qquad x \geq x_{min}, \qquad \alpha > 1$$

Exponential:

$$p_{exp}(x) = \lambda e^{-\lambda(x - x_{min})}, \qquad x \geq x_{min}, \qquad \lambda > 0$$

Lognormal:

$$p_{log}(x) = \frac{1}{x\sigma\sqrt{2\pi}} \frac{e^{\left(-\frac{(\ln(x)-\mu)^2}{2\sigma^2}\right)}}{\left(1 - \Phi\left(\frac{\ln(x_{min}) - \mu}{\sigma}\right)\right)}, \qquad x \geq x_{min}$$

where $\Phi(\cdot)$ denotes the cumulative distribution function of the standard normal distribution.

Across the candidate-model comparison, the lognormal distribution provided the most consistent fit for the empirical feature distributions across groups. Therefore, the lognormal model was used for subsequent modeling.

For each feature, the lognormal distribution was parameterized as $\ln(X) \sim \mathcal{N}(\mu, \sigma^2)$, where $\mu$ and $\sigma$ denote the mean and standard deviation of $\ln(X)$, respectively.

Supplementary Table S1 reports the parameters of the lognormal models fitted to the key features for each group (AD, PD, and HC). Parameter estimation was performed using the full (non-truncated) data.

Supplementary Table S1. Parameters of the lognormal models fitted to the SHAP-identified key features for AD, PD,and HC groups. Values are reported as $\mu$ ( $\sigma$) in log-space.

| **Feature** | **AD**<br>$\mu$ ( $\sigma$) | **PD**<br>$\mu$ ( $\sigma$) | **HC**<br>$\mu$ ( $\sigma$) |
|---|---|---|---|
| $\theta/\alpha$ ratio | -0.35 (0.79) | -0.01 (0.75) | -1.07 (0.48) |
| Mean relative $\theta$ power | -1.70 (0.41) | -1.54 (0.35) | -2.11 (0.36) |
